# Lamin A/C controls nuclear matrin-3 levels and localization, but not alternative splicing of cassette exons

**DOI:** 10.1101/378240

**Authors:** Dipen Rajgor, Clare Gooding, Robert Hayward, Miguel B Coelho, Christopher WJ Smith, Catherine M Shanahan

## Abstract

Disruptions in connections between the nuclear lamina and nuclear matrix occur in myopathic disorders. However, the biological significance of nuclear lamina - nuclear matrix coupling still remains largely undetermined. Previously it has been demonstrated that the nuclear matrix protein, matrin-3, binds to lamin A/C and this interaction is disrupted in laminopathies resulting in enhanced separation between the lamina and matrix. Matrin-3 has recently been identified as a core regulator of alternative splicing, whereas the involvement of lamin A/C in splicing still remains controversial. In this study, we demonstrate that lamin A/C is not only required for maintaining the nuclear organization of matrin-3, but also of other splicing activators and small nuclear ribonucleoproteins (snRNP) components. Interestingly, mis-localization of these splicing components did not appear to significantly disrupt alternative splicing events of cassette exons regulated by matrin-3. Thus, the lamin A/C-matrin3 interaction is unlikely to be involved in controlling alternative splicing but could be important in coordinating other nuclear activities. Interestingly, matrin-3 knock-down results in misshapen nuclei suggesting its interaction with lamin A/C maybe important in maintaining nuclear structural integrity.

## Introduction

The nuclear lamina underlies the nuclear face of the nuclear envelope and is composed of filamentous protein networks which provide structural support to the nucleus [1]. The lamina organizes sub-nuclear components such as chromatin, PML bodies and transcription factors [2]. The lamina is a core component of the Linker of Nucleoskeleton and Cytoskeleton (LINC) complex and therefore is integral in transducing signals between the nucleus and cytoskeleton [2]. Understanding the structural and functional properties of the nuclear lamina is necessary as mutations in lamina proteins cause a plethora of diseases known as laminopathies, which manifest as a broad range of tissue specific pathologies including different forms of premature ageing, muscular dystrophy, cardiomyopathy, lipodystrophy and neuropathy disorders [3]. At the cellular and molecular level, laminopathy mutations cause various perturbations in nuclear structure, integrity and positioning as well as defects in cell migration, mechanosensing and mechanotransduction [4–6].

The nuclear matrix is a protein-rich insoluble network that extends throughout the nucleoplasm and is classically defined as being resistant to high salt or detergent extraction [7, 8]. Multiple proteins involved in RNA processing, transcription and DNA replication are components of the nuclear matrix and it therefore plays critical roles in coordinating nuclear activity [8–10]. Matrin-3 is a multi-functional nuclear matrix protein with a wide range of roles in all of the above functions [11–14]. However, recent studies demonstrate matrin-3’s main nuclear role is likely to be in regulating alternative splicing of specific cassette exons and in repressing splicing of intronic LINE elements [14–16]. Matrin-3 mutations have been detected in patients suffering from amyotrophic lateral sclerosis (ALS) and congenital heart defects, but it is unclear which of its cellular functions are hampered in these pathological disorders [17–19].

To date, little is known about how the nuclear lamina is connected to the nuclear matrix and the mechanistic coupling between the 2 compartments remain elusive. Recently, matrin-3 has been shown to interact with lamin A/C and disruption of this interaction caused increased separation between lamin A/C in the lamina and matrin-3 in the matrix [20]. In this study, we further examine the functional relationship between lamin A/C and matrin-3. We show that lamin A/C knock-down results in a reduction of various splicing components including nuclear matrin-3. Importantly, lamin A/C knockdown does not significantly affect matrin-3 regulated alternative splicing events, suggesting lamin A/C - matrin-3 complexes are unlikely to be involved in regulating alternative splicing. However, we show that matrin-3 is required for maintaining nuclear shape which indicates matrin-3 is an important structural component for preserving nuclear morphology.

## Results and Discussion

### Nuclear matrin-3 localization is lamin A/C dependent

Lamin A/C has previously been shown to have a major impact on the organization of sub-nuclear compartments such as PML bodies and chromatin [2, 21]. Recently it has also been shown to interact with the nuclear matrix component matrin-3, and disruptions in this interaction have been identified in laminopathies [20]. To further characterize the relationship between lamin A/C and matrin-3, we examined nuclear matrin-3 organization in U2OS cells depleted of lamin A/C using siRNA-mediated knock-down. siLamin A/C consistently reduced lamin A/C levels by ~60% 72 hours post transfection as determined by immunofluorescence microscopy (Figure 1A,B). Unexpectedly, we also saw a ~40% reduction in nuclear matrin-3 in response to lamin A/C knock-down over the same time period (Figure 1C,D). In addition to cells showing a loss of nuclear matrin-3, a subset of cells had uneven nuclear matrin-3 localization rather than uniform nuclear distribution seen in control cells (Supplementary Figure 1). As we did not observe increased cytoplasmic matrin-3 staining in lamin A/C knock-down cells, lamin A/C may either stabilizes matrin-3 in the nucleus or regulates its expression at the transcriptional level. qRT-PCR showed no significant change in the amount of matrin-3 mRNA in lamin A/C knock-down cells compared to control cells (Supplementary Figure 2) suggesting that lamin A/C is required for nuclear stability and proper distribution of matrin-3.

**Figure 1.**
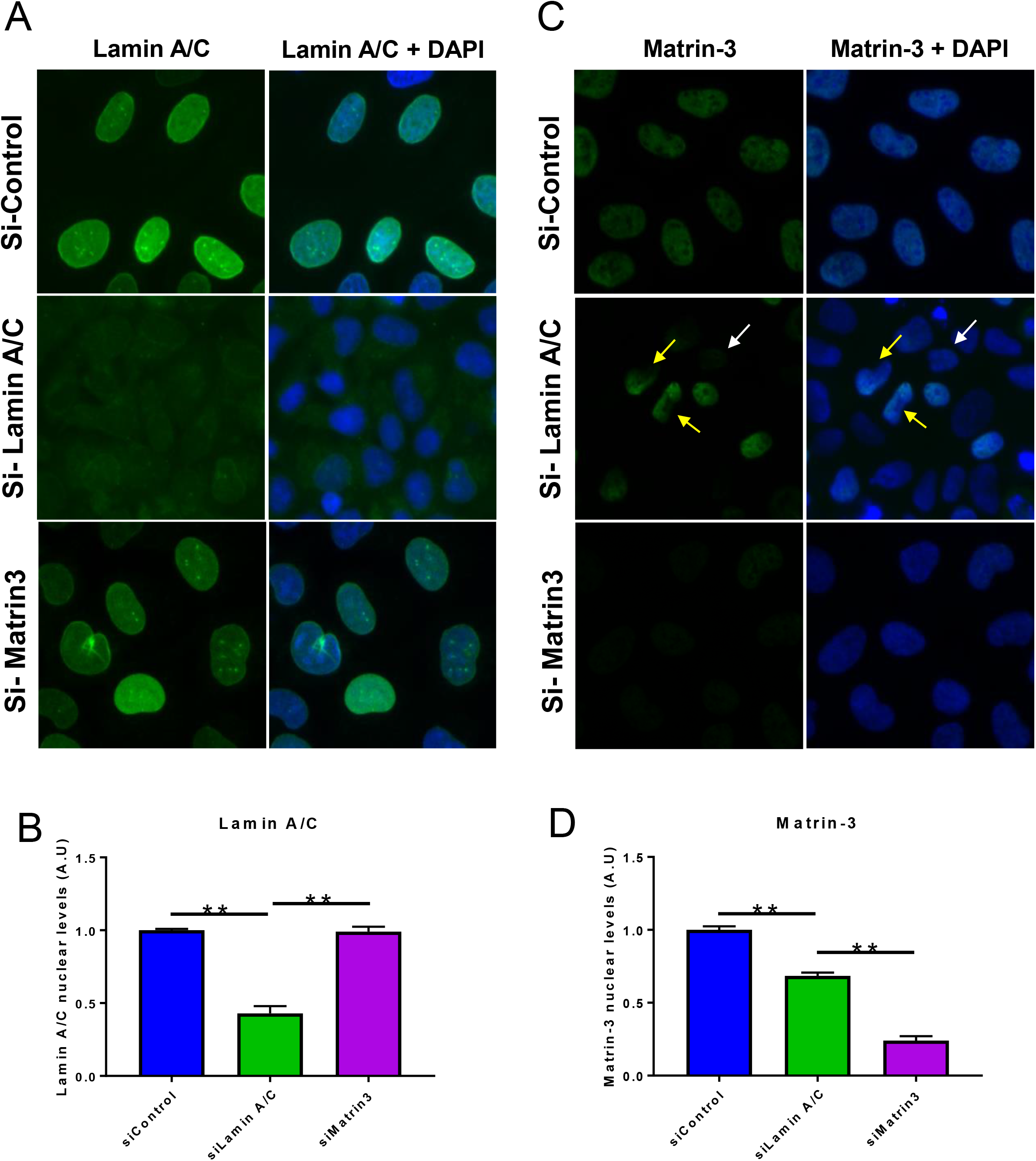
Nuclear Matrin-3 localization is dependent on lamin A/C. *A)* Lamin A/C staining in siRNA-mediated lamin A/C and matrin-3 knock-down U2OS cells. *B)* Quantification of nuclear lamin A/C levels in siRNA-mediated lamin A/C and matrin-3 knock-down U2OS cells. *C)* Matrin-3 staining in siRNA-mediated lamin A/C and matrin-3 knock-down U2OS cells. *D)* Quantification of nuclear matrin-3 levels in siRNA-mediated lamin A/C and matrin-3 knock-down U2OS cells. *N = >100 cells from 3 independent experiments. **P = <0.01 one-way ANOVA, Dunnett’s post hoc test.*

Next, we examined whether matrin-3 knock-down in U2OS affects the nuclear envelope distribution of lamin A/C. siMatrin-3 reduced matrin-3 expression by ~75% (Figure 1C,D), but did not significantly reduce the levels or distribution of lamin A/C (Figure 1A,B), indicating lamin A/C stability or localization is not matrin-3 dependent. Thus, matrin-3 stability and localization is dependent on lamin A/C, but not *vice versa.*

### Matrin-3 regulated alternative splicing events are largely lamin A/C independent

Next we examined the potential functional implications of matrin-3 mislocalizaiton due to nuclear lamina disruption and hypothesised it would also lead to changes in alternative splicing. To test this, we performed RT-PCR for cassette exons which were previously shown to be strongly regulated by matrin-3 in HeLa cells. We performed RT-PCR from cDNA isolated from matrin-3 depleted U2OS cells using primers in flanking constitutive exons and the percentage exon inclusion was determined (Figure 2). Matrin-3 knock-down in U2OS cells resulted in significant splicing changes in 5 of 6 cassette exons as previously seen in HeLa cells [14, 15]. Matrin-3 knock-down enhanced inclusion of cassette exons in *ST7, SETD5, PLEKHA3, PIGX* and *ACSL3*, while no change in splicing of exon 18 of *TCF12* was observed. In contrast, inclusion of *ST7* exon 11 showed a small (3.4%) but significant increase in response to lamin A/C knock-down. This effect was significantly less than the 10.8% increased exon inclusion in matrin-3 knock-down cells. None of the other 5 exons were affected by knock-down of lamin A/C despite showing increased inclusion of up to 45% upon matrin-3 knockdown (Figure 2). This suggests that the disruption of matrin-3 levels and localization caused by lamin A/C depletion is not sufficient to largely disrupt regulation of alternative splicing by matrin-3. Of note, although lamin A/C knock-down reduces nuclear matrin-3 levels, it is not to the same extent as those in cells transfected with siMatrin-3 (Figure 1D). Therefore it is plausible that sufficient levels of nuclear matrin-3 are present in lamin A/C knock-down cells to maintain matrin-3 regulated splicing events, or the nuclear pool of matrin-3 which is involved in splicing is not lost or mis-localized is response to lamin A/C knock-down. This would account for why matrin-3 regulated alternative splicing events are largely unaffected in lamin A/C depleted cells.

**Figure 2.**
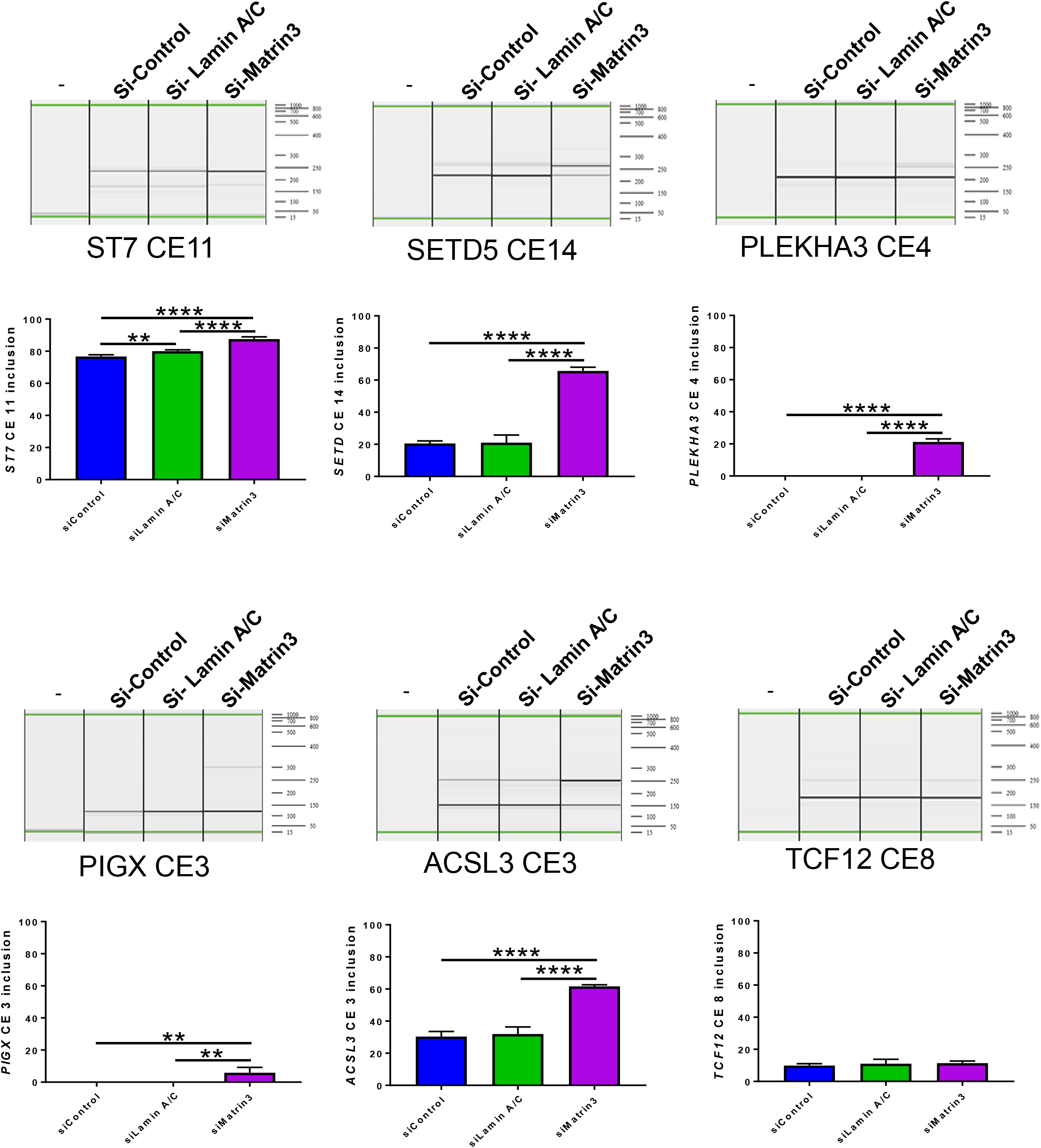
Lamin A/C is not largely involved in matrin-3 dependent cassette exon alternative splicing events. RT-PCR for matrin-3 regulated alternative cassette exon (CE) splicing events in the ST7, SETD5, PLEKHA3, PIGX, ACSL3 and TCF12 genes. Quantified percentage exon inclusion and standard deviations are plotted. *N = 5. **P = <0.01, ****P = <0.001 one-way ANOVA, Dunnett’s post hoc test.*

### Localization of other splicing factors are lamin A/C dependent

Whereas the nuclear lamina has been identified as a scaffold for key nuclear proteins such as transcription factors, no direct evidence exists for its involvement in scaffolding splicing components [2, 22]. However, a recent proteomic screen identified lamin A/C to interact with a number of splicing factors and spliceosomal components [20]. To determine if the nuclear localization of other splicing regulatory proteins were also dependent on lamin A/C, we examined a panel of splicing components using immunofluorescence microscopy on lamin A/C knock-down U2OS cells. Nuclear levels of the core small nuclear ribonucleoproteins (snRNP) components U2 snRNPA, U1-70K and SmD1 and the splicing factor SRSF6 all displayed up to 50% reduction in lamin A/C knock-down cells, similar to matrin-3 (Figure 3A,B,C,D). In contrast, the splicing activator SRSF1 and TRA2B maintained nuclear levels in these cells suggesting that an intact nuclear lamina may be required for maintaining levels of a subset of splicing factors within the nucleus (Figure 3E,F). SRSF1 localization being lamin A/C independent is consistent with previous findings [23]. Interestingly, we saw some cytoplasmic foci containing SRSF6 in lamin A/C knock-down cells, although these foci were not able to account fully for the loss of nuclear protein (Figure 3C). Likewise, SmD1 localization was more cytoplasmic in response to lamin A/C knock-down but did not form cytoplasmic foci (Figure 3D). In contrast, no cytoplasmic redistribution of U1-70K, U2 snRNPA, TRA2B and SRSF1 was observed in response to lamin A/C knock-down (Figure 3A,B,D,E,F).

**Figure 3.**
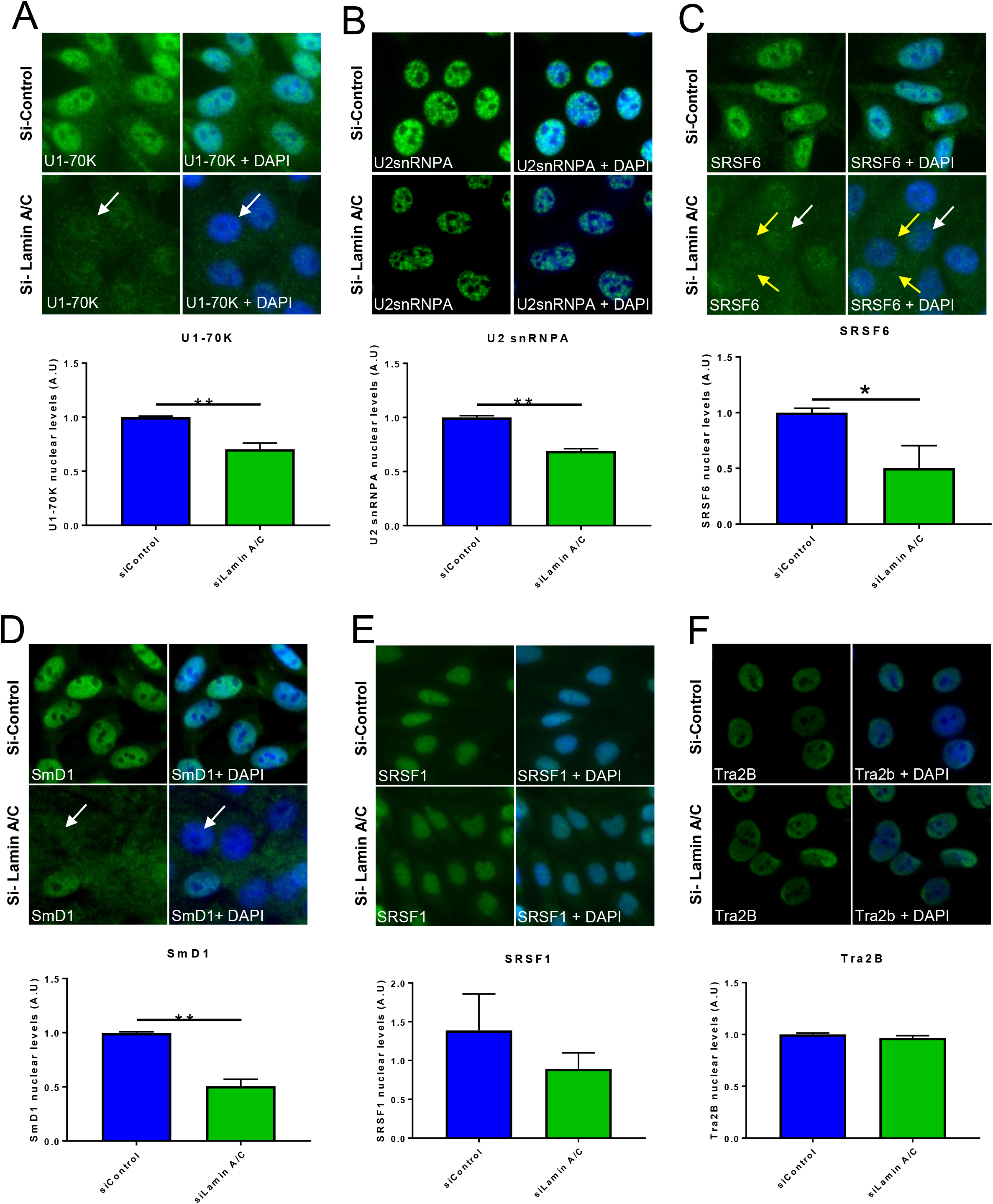
Nuclear U1-70K, U2 snRNP A, SRSF6, and smD1 localization is dependent on lamin A/C. siRNA-mediated lamin A/C knock-down results in a significant loss of nuclear *A)* U1-70K, *B)* U2 snRNP A, *C)* SRSF6 and *D)* SmD1, as determined by immunofluorescence microscopy. White arrowheads show example of nuclear loss of splicing components in lamin A/C knock-down cells. Yellow arrow heads show cytoplasmic foci in lamin A/C knock-down cells. *E)* SRSF1 and *F)* Tra2B nuclear levels are not significantly reduced in lamin A/C knock-down cells. *N = > 100 cells from 3 independent experiments. *P = < 0.05; **P = < 0.01; Students T-test.*

In this study we only examined the levels and localization of splicing components and alternative splicing events in U2OS cells depleted of lamin A/C, rather than diseased cells/tissue expressing *LMNA* mutations. Future studies should focus on expressing *LMNA* mutants, for example *LMNA* Δ303 which is unable to interact with matrin-3, into relevant cell lines and using robust sequencing methods, such as mRNA-seq, to obtain a more physiological relevant read out of lamin A/C regulated mRNA cassette exon splicing. These experiments will truly provide key information into whether mis-localization of splicing factors or disruptions in alternative splicing are casual of laminopathies

### Matrin-3 is required for maintaining nuclear shape, but not nuclear size

As lamin A/C depletion did not alter matrin-3 regulated alternative splicing events, we assessed whether matrin-3 could be involved in regulating structural aspects of the nucleus. The reduction of lamin A/C or any components of the LINC complex are known to promote nuclear morphology defects [24]. As matrin-3 interacts with lamin A/C and it’s nuclear localization is lamin A/C dependent, we also analysed nuclear morphology based on nuclear circularity in lamin A/C and matrin-3 knock-down cells. As expected, lamin A/C depleted cells were less circular than control cells and matrin-3 knock-down cells. Although matrin-3 depleted cells were significantly less mis-shapen than lamin A/C depleted cells, they were not as circular as control cells (Figure 4A). Lamin A/C knock-down has also been reported to reduce nuclear size [25]. In our experiments, we observed a reduction in nuclear size by ~20% in response to lamin A/C knock-down. However, nuclei in matrin-3 knock-down cells were similar in size to control cells (Figure 4B). Together, this data suggests matrin-3 and lamin A/C may be required for maintaining nuclear shape, but not nuclear area, and it is possible that the nuclear matrin-3 loss / mis-localization in lamin A/C depleted cells contributes to the grossly mis-shapen nuceli seen in these cells and in laminopathies.

**Figure 4.**
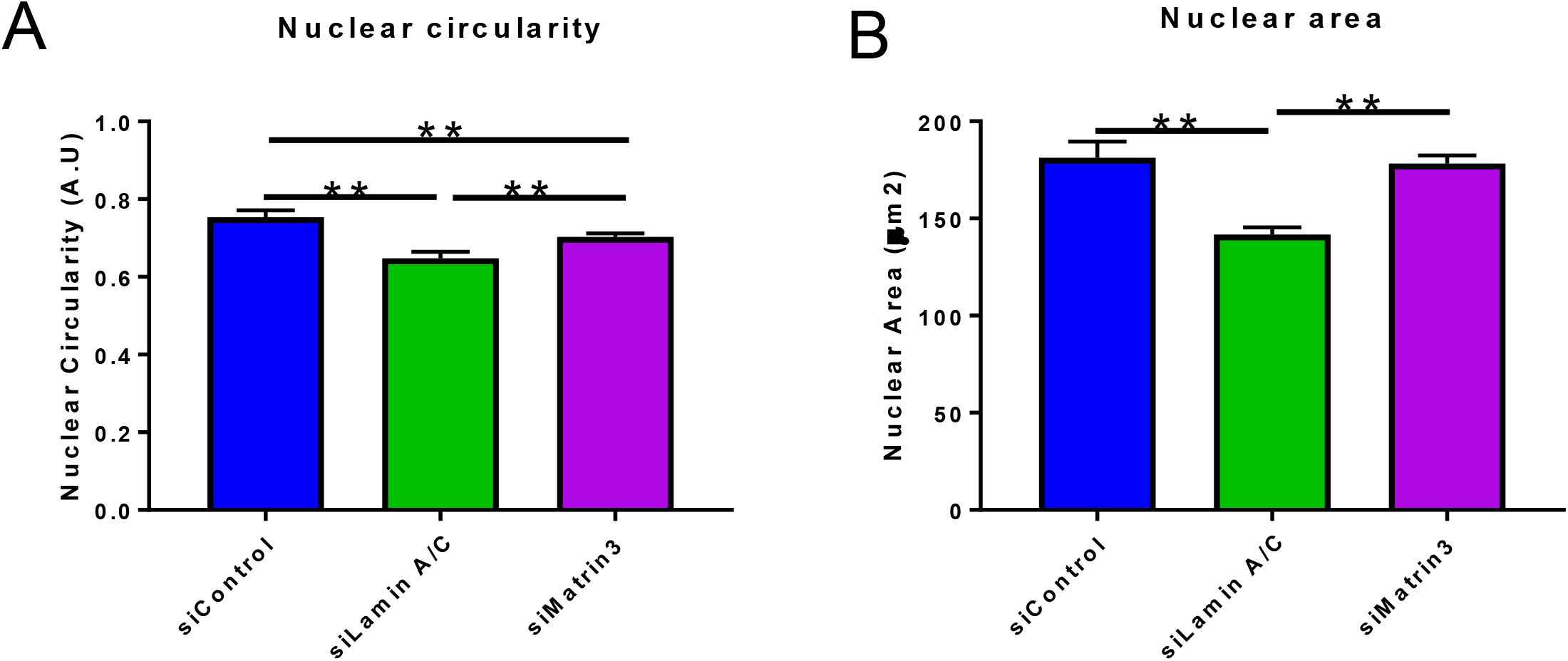
Matrin-3 regulates nuclear shape, but not nuclear size. *A)* Lamin A/C and matrin-3 knock-down result in mis-shapen nuclei based on nuclei circularity measurements. *B)* Lamin A/C knock-down cells, but not matrin-3 knock-down cells, have smaller nuclei. *N = >100 cells from 3 independent experiments. **P = <0.01 one-way ANOVA, Dunnett’s post hoc test.*

### Conclusions

Matrin-3 is a multi-functional protein which has been implicated in a range of nuclear events and lamin A/C is a major nuclear scaffold important for a host of nuclear activities. Therefore, it is likely that matrin-3-lamin A/C complexes have additional functional roles in addition to maintaining nuclear structural shape. Unrelated studies indicate matrin-3 and lamin A/C have some overlapping functionality. For example, both matrin-3 and lamin A/C interact with chromatin binding proteins and various transcription factors, suggesting they could play important gene regulatory roles [11, 22]. Interestingly, both proteins are involved in triggering DNA damage responses induced by DNA double stranded breaks [26, 27]. Both matrin-3 and lamin A/C interact with KU70, which binds to broken DNA ends and recruits repair enzymes to facilitate repair [26, 28]. Knock-down of either matrin-3 or lamin A/C significantly delays the recruitment time of repair factors to regions of DNA damage, suggesting the matrin-3 and lamin A/C interaction may be important in repairing DNA damage [26, 27]. Importantly, DNA damage occurs in laminopathies and this may be due to disrupted lamin A/C-matrin-3 scaffolds at DNA damaged sites. Therefore future studies should focus on identifying additional roles for lamin A/C-matrin-3 complexes as this is likely to provide mechanistic insight into the molecular defects which occur in laminopathies and diseases associated with matrin-3 mutations.

## Materials and Methods

### Cell Culture

U2OS cells were passaged when reaching ~70% confluency and maintained in DMEM complete media (Sigma) supplemented with 10U/ml penicillin, 10mg/ml streptomycin, 10mg/ml Glutamine and 10% foetal bovine serum.

### Immunofluorescence microscopy

Cells were fixed for 5 min in 4% PFA (Sigma-Aldrich) followed by 2 min permeabilization with 0.5% NP-40 (EMD Millipore). The coverslips were incubated with blocking solution (1% BSA) for 1 hour at room temperature. Primary antibodies used for immuno-staining were diluted in blocking solution and applied to the coverslips for 1 hour at room temperature followed by three short rinses in PBS. Primary antibodies were as follows: anti-Matrin-3 (Abcam), anti-Lamin A/C (Santa Cruz), anti-SRSF6 (Santa Cruz), anti-U2 snRNPA (Santa Cruz), anti-U1-70K (Millipore) or anti-smD1 (Santa Cruz). Next, the coverslips were incubated with secondary antibodies, diluted 1:1,000 in blocking buffer, conjugated to Alexa Fluor 488 (Invitrogen) for 1 hour at room temperature. The coverslips were next stained with DAPI and rinsed in PBS before being mounted onto slides using Mowiol mounting media. Images were acquired at room temperature using a using an Olympus IX81 wide-field microscope with a 40x air objective lens attached to a Hamamatsu Orca-R2 cooled CCD camera.

### Image analysis

Nuclear levels of all proteins were measured by fluorescence integrated density in ImageJ. Nuclear circularity measurements were taken using the ‘Analyse particles’ function in ImageJ on thresholded DAPI staining. Circularity values were given between 0 and 1, with values closer to 1 being more circular [29, 30]. All experiments were repeated at least 3 or more times and between 100-150 cells were analyzed. The mean readings per experiment were used for statistical analysis in GraphPad Prism. Student’s T-test or one-way Annova were performed as indicated in the figure legends.

### qRT-PCR

Total RNA was isolated from U2OS cells using Triazole RNA STAT-60 and phenol chloroform extraction [31]. 2 ug of total RNA was reverse transcribed using AMV Reverse Transcriptase (Promega) according to the manufacturer’s instructions. qPCR was performed in a 20 ul reaction containing cDNA per 1X SYBR Green PCR master mix (Eurogentec) and 0.1 mM of each primer. Matrin-3 forward (5’TCGTCGATGCCAGCTTCTTCTTGA3’), matrin-3 reverse (5’C CTTG C AG GTTT C C ATTT C CAG C A3’), GAPDH forward (5’CGACCACTTTGTCAAGCTC3’) and GAPDH reverse (5’CAAGGGTCTACATGGCAAC3’) primers were used. The cycling parameters were 94°C for 15 seconds followed by a single step annealing and extension at 60°C for 60 seconds. Amplifications were performed on RotorGene-3000 (Corbett). Fold changes between samples were calculated by the delta-delta CT method.

### RT-PCR for cassette exon splicing

RNA was isolated from U2OS cells and reverse transcribed to cDNA as described above. PCRs for cassette exon splicing were carried out using primer sets in flanking constitutive exons as previously described [14, 15]. All PCRs were carried out using the Jumpstart Taq polymerase (Sigma), and the products were separated and quantified on a QIAXcel capillary electrophoresis system (Qiagen) as previously described[14].

## Acknowledgments

This work was funded by a Medical Research Council (MRC) centenary fellowship award to DR, Wellcome Trust programme grant (092900) awarded to CWJS and a British Heart Foundation (BHF) programme grant awarded to CMS.

## References

1. Gruenbaum, Y., et al., The nuclear lamina comes of age. Nat Rev Mol Cell Biol, 2005. 6(1): p. 21–31.

2. Dechat, T., et al., Nuclear lamins: major factors in the structural organization and function of the nucleus and chromatin. Genes Dev, 2008. 22(7): p. 832–53.

3. Worman, H.J. and G. Bonne, “Laminopathies”: a wide spectrum of human diseases. Exp Cell Res, 2007. 313(10): p. 2121–33.

4. O’Leary, M.N., et al., The ribosomal protein Rpl22 controls ribosome composition by directly repressing expression of its own paralog, Rpl22l1. PLoS Genet, 2013. 9(8): p. e1003708.

5. Lee, J.S., et al., Nuclear lamin A/C deficiency induces defects in cell mechanics, polarization, and migration. Biophys J, 2007. 93(7): p. 2542–52.

6. Lammerding, J., et al., Lamin A/C deficiency causes defective nuclear mechanics and mechanotransduction. J Clin Invest, 2004. 113(3): p. 370–8.

7. Verheijen, R., W. van Venrooij, and F. Ramaekers, The nuclear matrix: structure and composition. J Cell Sci, 1988. 90 (Pt 1): p. 11–36.

8. Razin, S.V., et al., Nuclear matrix and structural and functional compartmentalization of the eucaryotic cell nucleus. Biochemistry (Mosc), 2014. 79(7): p. 608–18.

9. Nickerson, J., Experimental observations of a nuclear matrix. J Cell Sci, 2001. 114(Pt 3): p. 463–74.

10. Skowronska-Krawczyk, D. and M.G. Rosenfeld, Nuclear matrix revisited? Cell Cycle, 2015. 14(10): p. 1487–8.

11. Zeitz, M.J., et al., Matrin 3: chromosomal distribution and protein interactions. J Cell Biochem, 2009. 108(1): p. 125–33.

12. Zhang, Z. and G.G. Carmichael, The fate of dsRNA in the nucleus: a p54(nrb)-containing complex mediates the nuclear retention of promiscuously A-to-I edited RNAs. Cell, 2001. 106(4): p. 465–75.

13. Salton, M., et al., Involvement of matrin 3 and SFPQ/NONO in the DNA damage response. Cell Cycle, 2010. 9(8).

14. Coelho, M.B., et al., Nuclear matrix protein Matrin3 regulates alternative splicing and forms overlapping regulatory networks with PTB. EMBO J, 2015. 34(5): p. 653–68.

15. Uemura, Y., et al., Matrin3 binds directly to intronic pyrimidine-rich sequences and controls alternative splicing. Genes Cells, 2017. 22(9): p. 785–798.

16. Jan Attig, F.A., Clare Gooding, Aarti Singh, Anob M, Chakrabarti, Nejc Haberman, Warren Emmett, Christopher WJ Smith, Nicholas M Luscombe and Jernej Ule, Heteromeric RNP assembly at LINEs controls lineage-specific RNA processing. BioRxiv, 2018.

17. Johnson, J.O., et al., Mutations in the Matrin 3 gene cause familial amyotrophic lateral sclerosis. Nat Neurosci, 2014. 17(5): p. 664–666.

18. Marangi, G., et al., Matrin 3 variants are frequent in Italian ALS patients. Neurobiol Aging, 2017. 49: p. 218 e1–218 e7.

19. Quintero-Rivera, F., et al., MATR3 disruption in human and mouse associated with bicuspid aortic valve, aortic coarctation and patent ductus arteriosus. Hum Mol Genet, 2015. 24(8): p. 2375–89.

20. Depreux, F.F., et al., Disruption of the lamin A and matrin-3 interaction by myopathic LMNA mutations. Hum Mol Genet, 2015. 24(15): p. 4284–95.

21. Houben, F., et al., Cytoplasmic localization of PML particles in laminopathies. Histochem Cell Biol, 2012.

22. Andres, V. and J.M. Gonzalez, Role of A-type lamins in signaling, transcription, and chromatin organization. J Cell Biol, 2009. 187(7): p. 945–57.

23. Vecerova, J., et al., Formation of nuclear splicing factor compartments is independent of lamins A/C. Mol Biol Cell, 2004. 15(11): p. 4904–10.

24. Polychronidou, M. and J. Grobhans, Determining nuclear shape: the role of farnesylated nuclear membrane proteins. Nucleus, 2011. 2(1): p. 17–23.

25. Jevtic, P., et al., Concentration-dependent Effects of Nuclear Lamins on Nuclear Size in Xenopus and Mammalian Cells. J Biol Chem, 2015. 290(46): p. 27557–71.

26. Salton, M., et al., Involvement of Matrin 3 and SFPQ/NONO in the DNA damage response. Cell Cycle, 2010. 9(8): p. 1568–76.

27. Singh, M., et al., Lamin A/C depletion enhances DNA damage-induced stalled replication fork arrest. Mol Cell Biol, 2013. 33(6): p. 1210–22.

28. Kubben, N., et al., Identification of differential protein interactors of lamin A and progerin. Nucleus, 2010. 1(6): p. 513–25.

29. Wheeler, M.A., et al., Identification of an emerin-beta-catenin complex in the heart important for intercalated disc architecture and beta-catenin localisation. Cell Mol Life Sci, 2010. 67(5): p. 781–96.

30. Zhou, C., et al., Novel nesprin-1 mutations associated with dilated cardiomyopathy cause nuclear envelope disruption and defects in myogenesis. Hum Mol Genet, 2017. 26(12): p. 2258–2276.

31. Rajgor, D., et al., Multiple novel nesprin-1 and nesprin-2 variants act as versatile tissue-specific intracellular scaffolds. PLoS One, 2012. 7(7): p. e40098.

